# Improved conditioning for hematopoietic chimerism induces islet tolerance to cure diabetes

**DOI:** 10.1101/2025.06.26.661773

**Authors:** Stephan A. Ramos, Preksha Bhagchandani, Diego M. Burgos, Xueying Gu, Richard Rodriguez, Nadia Nourin, Martin Neukam, Shiva Pathak, Judith A. Shizuru, Seung K. Kim

## Abstract

Mixed hematopoietic chimerism after hematopoietic cell transplantation (HCT) can modulate the immune system and induce tolerance to allogeneic tissues. However, bone marrow conditioning-related toxicities preclude wider adoption of HCT for transplant allotolerance. We sought agents that reduced conditioning intensity, while promoting durable mixed chimerism after HCT across complete major histocompatibility complex (MHC) mismatch in diabetic mice, permitting islet allotransplantation and diabetes reversal. We systematically tested baricitinib (JAK1/2 inhibitor), venetoclax (Bcl2 inhibitor), and αCD47 antibody, agents in current clinical use, and quantified hematopoietic chimerism after HCT. Combined with αCD117 antibody, transient T cell depletion, and just 10 centigray (cGy) total body irradiation (TBI), these agents enabled durable mixed chimerism and matching allo-islet tolerance, to cure diabetes without evidence of GVHD. Thus, we have developed a conditioning regimen to promote allogeneic mixed hematopoietic chimerism and transplanted islet allotolerance that minimizes conditioning radiation and cures diabetes, a significant achievement.

## Introduction

Transplantation of MHC-mismatched allogeneic islets is an FDA-approved approach to reverse diabetes after pancreatic β cell loss (1,2). Though effective for restoring glycemic control, standard clinical practice uses chronic immunosuppression to prevent islet allorejection (3), which is associated with significant toxicities. These include β cell and renal injury, increased risk of opportunistic infections and malignancies, and chronic rejection, morbidities that have limited widespread adoption of islet allotransplantation in diabetes (4). Thus, islet cell replacement strategies have focused intensively on establishing immune allotolerance without systemic immunosuppression.

Hematopoietic stem cells (HSCs) are multipotent stem cells that reside in specialized bone marrow niches and are capable of generating all blood and immune cell lineages (5–7). After HCT, complete hematopoietic and immune reconstitution by donor HSCs can cure life-threatening conditions like blood disorders and malignancies and can correct autoimmunity. Mixed hematopoietic chimerism after HCT, where recipient- and donor-derived HSCs co-exist, is an effective method to modulate the recipient immune system and establish tolerance to allogeneic donor-matched tissues and solid organs (8–12). Mixed chimerism can reshape central and peripheral immune tolerance, promoting deletion or suppression of allo- (donor) reactive and auto- (self) reactive immune cells while fostering regulatory immune cell populations that support acceptance of donor-matched allogeneic grafts. Beyond enabling immune tolerance, mixed chimerism can contribute to the suppression of autoreactive immune responses, offering potential therapeutic benefits for autoimmune diseases, like type 1 diabetes (13).

To achieve successful engraftment of donor HSCs and establish mixed hematopoietic chimerism, recipients undergo prior bone marrow ‘conditioning’. Conditioning is a preparative regimen that creates space in the bone marrow niche for donor HSC engraftment and transiently suppresses the recipient immune system to prevent acute graft rejection (14). However, current conditioning regimes were largely developed for malignancies and rely on complete or near-complete eradication of the recipient hematopoietic system (’myeloablative’) with radiation therapy (XRT) and/or genotoxic or cytotoxic chemotherapy. Thus, standard conditioning is associated with severe toxicities, including infections, secondary malignancies, and organ damage, preventing broader adoption of mixed chimerism-based approaches for non-malignant conditions such as diabetes (15,16). Mere dose reduction of myeloablative conditioning agents to mitigate conditioning intensity and toxicity unfortunately results in HSC engraftment failure (17). Although multiple agents have been explored to reduce conditioning intensity in preclinical models, clinical translation has been limited due to systemic toxicity, graft-versus-host-disease (GVHD), and inefficacy in humans (13,18–21). Thus, the development of safer non-myeloablative (NMA) conditioning regimes could advance and expand use of mixed hematopoietic chimerism for transplant tolerance (22).

Previously, we reported a NMA conditioning strategy incorporating anti-CD117 antibody and a single 300 cGy dose of TBI for curing established diabetes after allogeneic HCT and islet transplantation (23). However, this TBI dose is associated with increased risk for secondary malignancy and infertility, precluding its use in patient subsets, including children and women of childbearing age (23,24).

A promising strategy to achieve effective, less toxic conditioning involves targeting specific cellular pathways involved in immune regulation and hematopoietic niche remodeling (25,26). Baricitinib, a Janus kinase (JAK1/2) inhibitor, modulates immune responses by reducing inflammatory cytokine signaling and attenuating T and NK cell responses and enhancing engraftment (26,27). CD47 is a transmembrane protein expressed on many cell types, including HSCs, that interacts with SIRPα on neutrophils and macrophages to prevent antibody-dependent cell-mediated cytotoxicity and phagocytosis (28,29). Disrupting this “don’t eat me” signal on HSCs with αCD47 antibody therapy promotes the clearance of HSCs, and facilitates clearance of the bone marrow HSC niche (30,31). Venetoclax, is a BCL-2 inhibitor that selectively induces apoptosis in hematopoietic and immune cells while sparing non-target tissues and promotes donor HSC engraftment with reduced XRT (32). These clinically portable, non-genotoxic, agents act on distinct pathways to support donor cell engraftment while reducing conditioning toxicity. Here, when combined with αCD117 antibody, transient T cell depletion, and only 10 cGy TBI, these agents enabled durable mixed hematopoietic chimerism and matching allo-islet tolerance to cure established diabetes without evidence of GVHD. Thus, we present a novel, clinically relevant, NMA conditioning regimen to promote allogeneic mixed hematopoietic chimerism and transplanted islet allotolerance for diabetes.

## Results

### JAK1/2 inhibition permits reduced radiation in hematopoietic conditioning to achieve durable mixed chimerism

A prior NMA conditioning regimen we developed uses αCD117 antibody, transient T cell depletion (hereafter, ‘TCD’) with αCD4 and αCD8 antibodies, and 200-300 cGy TBI, to establish durable mixed hematopoietic chimerism after HCT with MHC-mismatched bone marrow (23). To improve conditioning further, we sought reagents with established clinical safety that would permit reduced radiation dosage. Baricitinib is an immune suppressive agent that inhibits NK and T cells via JAK1/2 blockade, and is well-tolerated in studies of allogeneic HSC engraftment (27,33). To evaluate if baricitinib could reduce conditioning TBI intensity, non-diabetic B6 CD45.1^+^ mice were administered baricitinib (400 µg/d) from days −6 to day +3, TCD from day −2 to day 0, irradiated with de-escalated doses of 75, 50 or 25 cGy TBI on day −3, transplanted with BALB/c CD45.2^+^ lineage-depleted hematopoietic stem and progenitor cells (HSPCs), and followed for up to 26 weeks (**Figure 1, A** and **B**). Four weeks after HCT, we observed donor engraftment in all peripheral blood lineages in mice conditioned with 75 cGy (**Figure 1C**). Stable mixed chimerism was maintained in blood throughout the 27-week experiment period in 5 of 5 mixed chimeric recipients (hereafter, “BALB/c:B6 mice”; **Figure 1D**). Endpoint analysis of the spleen and bone marrow showed comparable chimerism levels (**Figure 1, E** and **F**). We also confirmed bone marrow engraftment of *donor* Lin–Sca1+cKit+ (LSK) HSCs, and persistence of *host* LSK HSCs (donor LSK chimerism: 34.2 ± 25.4%; **Figure 1F**). By contrast, we did not achieve durable multilineage mixed chimerism in B6 mice after conditioning with 50 or 25 cGy (**Supplementary Figure 1, A-D**). Specifically, B6 mice conditioned with 50 or 25 cGy TBI did not generate donor-derived CD3^+^ T cells, which are crucial for allogeneic tolerance and stable mixed hematopoietic chimerism (34) (**Supplementary Figure 1, A-D**).

**Figure 1:**
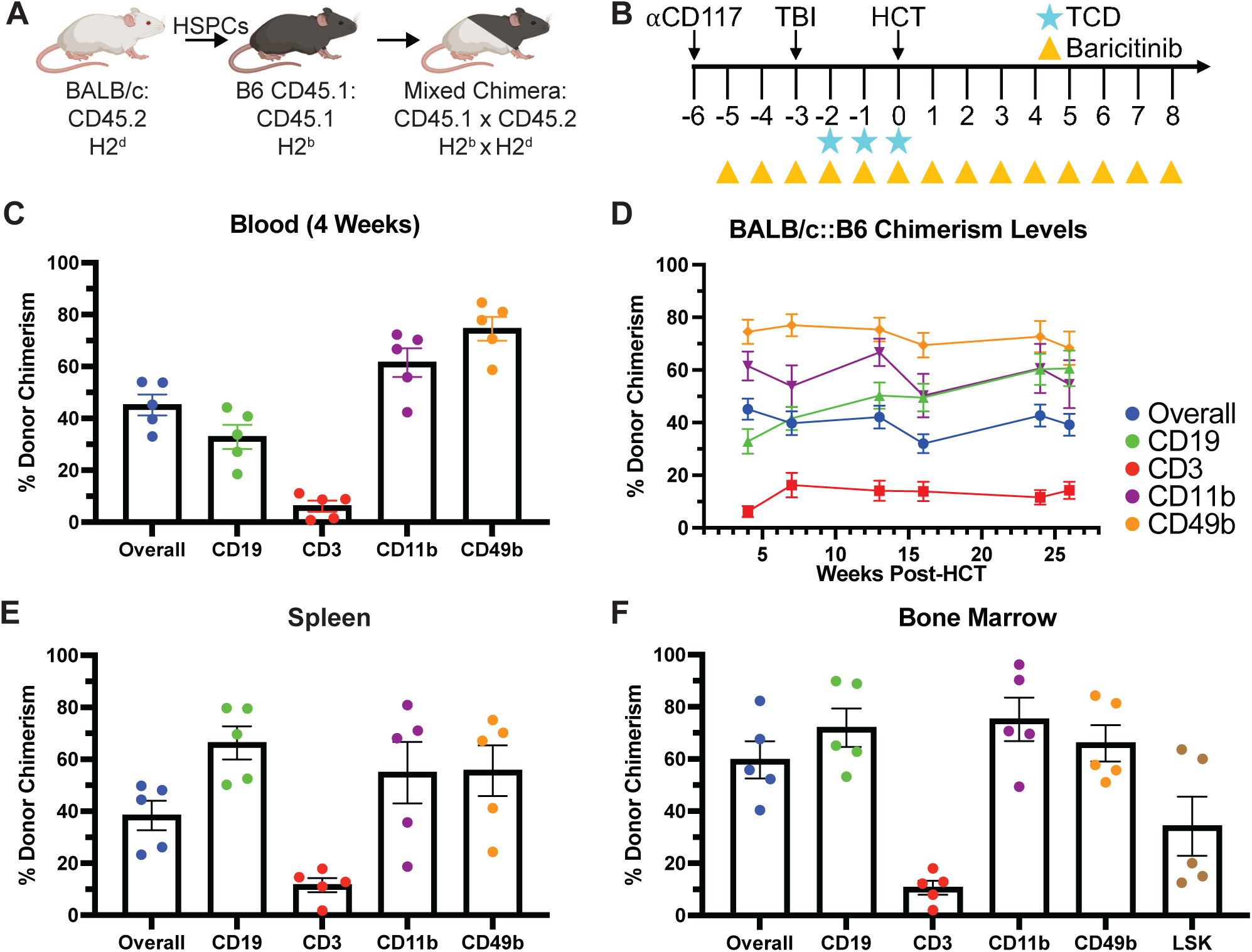
Non-myeloablative conditioning with baricitinib and 75 cGy TBI permits durable allogeneic donor chimerism. **(A)** Transplantation model and strains used. **(B)** Experimental conditioning and transplantation timeline. **(C)** Multilineage chimerism analysis of peripheral blood 4 weeks post-HCT. **(D)** Longitudinal multilineage chimerism analysis of peripheral blood through 26 weeks post-HCT. **(E)** Multilineage chimerism analysis of host spleen 26 weeks post-HCT. **(F)** Multilineage chimerism analysis, including Lin^-^Sca1^+^cKit^+^ (LSK) HSCs, of host bone marrow 26 weeks post-HCT. (C-F) n = 5. Data are represented with mean ± SEM. TBI = total body irradiation; HCT = hematopoietic cell transplant; TCD = T cell depletion. See also **Supplementary Figure 1**.

Using whole bone marrow (WBM) instead of lineage-depleted HSPCs achieved similar outcomes (**Supplementary Figure 2**). After HSPC or WBM transplantation, we did not observe signs of GVHD (e.g., mucosal changes, skin rash, or weight loss), and longitudinal body weight measures showed excellent weight gain from HCT to experimental endpoint in all groups (**Supplementary Figure 1E** and **2C**). Thus, we established durable mixed hematopoietic chimerism in non-diabetic B6 mice across complete MHC mismatch with a NMA regimen combining 75cGy TBI with αCD117, TCD, and baricitinib. These studies illustrate that systematic addition of reagents or steps could be useful for reducing TBI and conditioning toxicity, and we used this strategy to reduce TBI further. In all subsequent studies, we transplanted WBM, to model the current clinical standard.

### Studies of bone marrow clearance after conditioning with αCD47 antibody and Bcl2 inhibitor

Prior studies have incorporated antibodies to CD47 for conditioning and HCT that achieved mixed chimerism across a haplo-mismatched MHC barrier (31,35). To evaluate conditioning with αCD47 to promote mixed chimerism across a full MHC mismatch, we conditioned B6 CD45.1^+^ mice with αCD117, baricitinib, TCD, and αCD47 antibody (300 µg/d) from day −6 to day −2. On day −3 mice were irradiated with 25 cGy TBI and transplanted with 30e6 WBM cells from CD45.2^+^ BALB/c donors on day 0 (**Figure 2A**). Serial analysis of peripheral blood for up to 28 weeks after HCT revealed successful multilineage chimerism levels in B6 mice after conditioning with αCD47 antibody and 25 cGy TBI (**Figure 2, B** and **C**). Endpoint analysis of the spleen and bone marrow showed durable multilineage chimerism levels and engraftment of donor CD45.2^+^ LSK HSCs (Donor LSK chimerism: 47.4 ± 7.9%; **Figure 2, D** and **E**). We did not observe indices of GVHD in BALB/c:B6 chimeras throughout our studies. However, addition of conditioning αCD47 antibody resulted in ∼20% reduction in body mass, necessitating daily oral feeding supplementation for 7 days during and after conditioning (**Figure 2F**).

**Figure 2:**
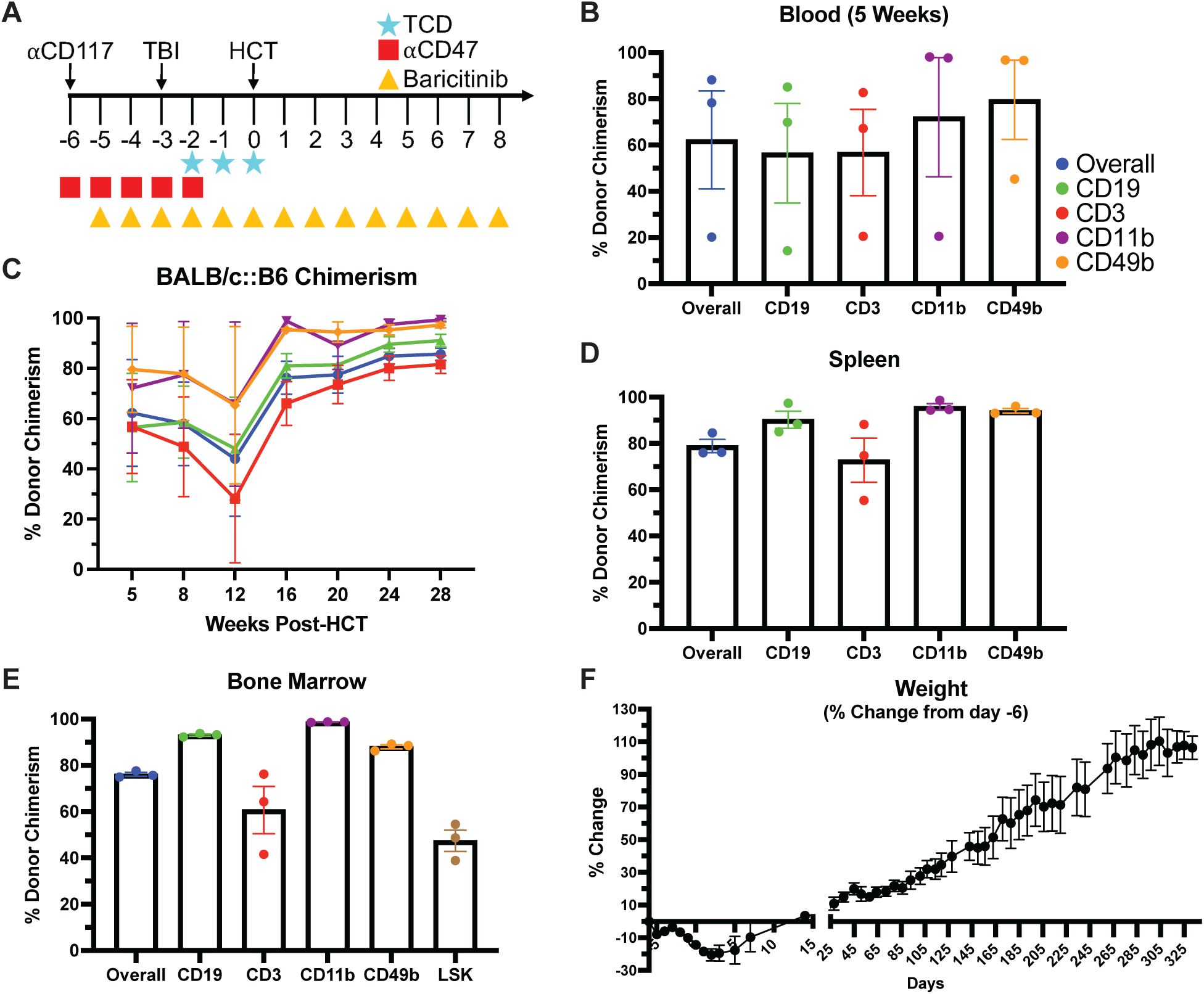
Anti-CD47 monoclonal antibody conditioning promotes durable mixed hematopoietic chimerism with 25 cGy TBI. **(A)** Experimental conditioning and transplantation timeline. **(B)** Multilineage chimerism analysis of peripheral blood 5 weeks post-HCT. **(C)** Longitudinal multilineage chimerism analysis of peripheral blood through 28 weeks post-HCT. **(D)** Multilineage chimerism analysis of host spleen 47 weeks post-HCT. **(E)** Multilineage chimerism analysis, including Lin^-^Sca1^+^cKit^+^ (LSK) HSCs, of host bone marrow 47 weeks post-HCT. **(F)** Weight after HCT of BALB/c:B6 as a percentage of initial weight prior to conditioning start. (B-E) n = 3. Data are represented with mean ± SEM. TBI = total body irradiation; HCT = hematopoietic cell transplant; TCD = T cell depletion.

To reduce conditioning toxicity, we sought αCD47 dosage reductions. We measured bone marrow clearance after conditioning with αCD117, baricitinib, 30 cGy TBI, and either 75 or 50 µg/d αCD47 (**Supplementary Figure 3A**). Conditioning with 75 and 50 µg/d αCD47 antibody for 5 consecutive days reduced overall bone marrow cellularity and HSC count equally (**Supplementary Figure 3, B** and **C**). We observed little to no change in body mass of mice conditioned with 50 µg/d, while the 75 µg/d group experienced significant weight loss (**Supplementary Figure 3D**). However, HSCs were not reduced to levels permissive to successful HCT engraftment with either αCD47 dose (**Supplementary Figure 3C**). These findings suggested that additional steps might be required with low dose αCD47 to permit successful HCT engraftment and further reduction of conditioning TBI dosing.

Venetoclax is a pharmacologic inhibitor of Bcl2, a survival factor in HSPCs and differentiated blood lineages, and was recently used to reduce conditioning TBI by 50% in non-human primates prior to successful mixed chimerism (26,32). We evaluated the impact of venetoclax (200 µg/d from days −6 to −2) on bone marrow clearance in mice conditioned with αCD117, baricitinib, and either 50, 25, or 10 cGy TBI (**Supplementary Figure 3E**).

Addition of venetoclax did not adequately clear HSCs, despite reduced overall cell counts (**Supplementary Figure 3, F** and **G**). However, venetoclax was well tolerated, causing only a mild, transient change in body mass that resolved by day 0 (**Supplementary Figure 3H**). We next investigated bone marrow clearance and mixed chimerism using αCD117-based conditioning that added both venetoclax and αCD47, a combination not been previously reported (26,32).

### Durable allogeneic mixed hematopoietic chimerism after conditioning with 10 cGy TBI

We conditioned B6 mice with venetoclax, αCD47 (50 µg/d), αCD117 antibody, TCD, baricitinib, and 10 cGy TBI (**Figure 3A**). This conditioning regimen led to durable multilineage mixed hematopoietic chimerism after BALB/c WBM transplantation (**Figure 3, B** and **C**), with only modest, transient weight loss (<10%) (**Figure 3F**). Endpoint analysis of blood, spleen, and bone marrow 20 weeks after HCT revealed mixed chimerism and engraftment of donor CD45.2^+^ LSK HSCs in the bone marrow (**Figure 3, D** and **E**). By contrast, mice failed to achieve multilineage mixed chimerism after omission of αCD47 antibody or TBI (0 cGy) from this regimen (**Supplementary Figure 4, A** and **B**). Thus, systematic testing identified a new combination of reagents that minimized conditioning XRT and permitted mixed chimerism in non-diabetic mice.

**Figure 3:**
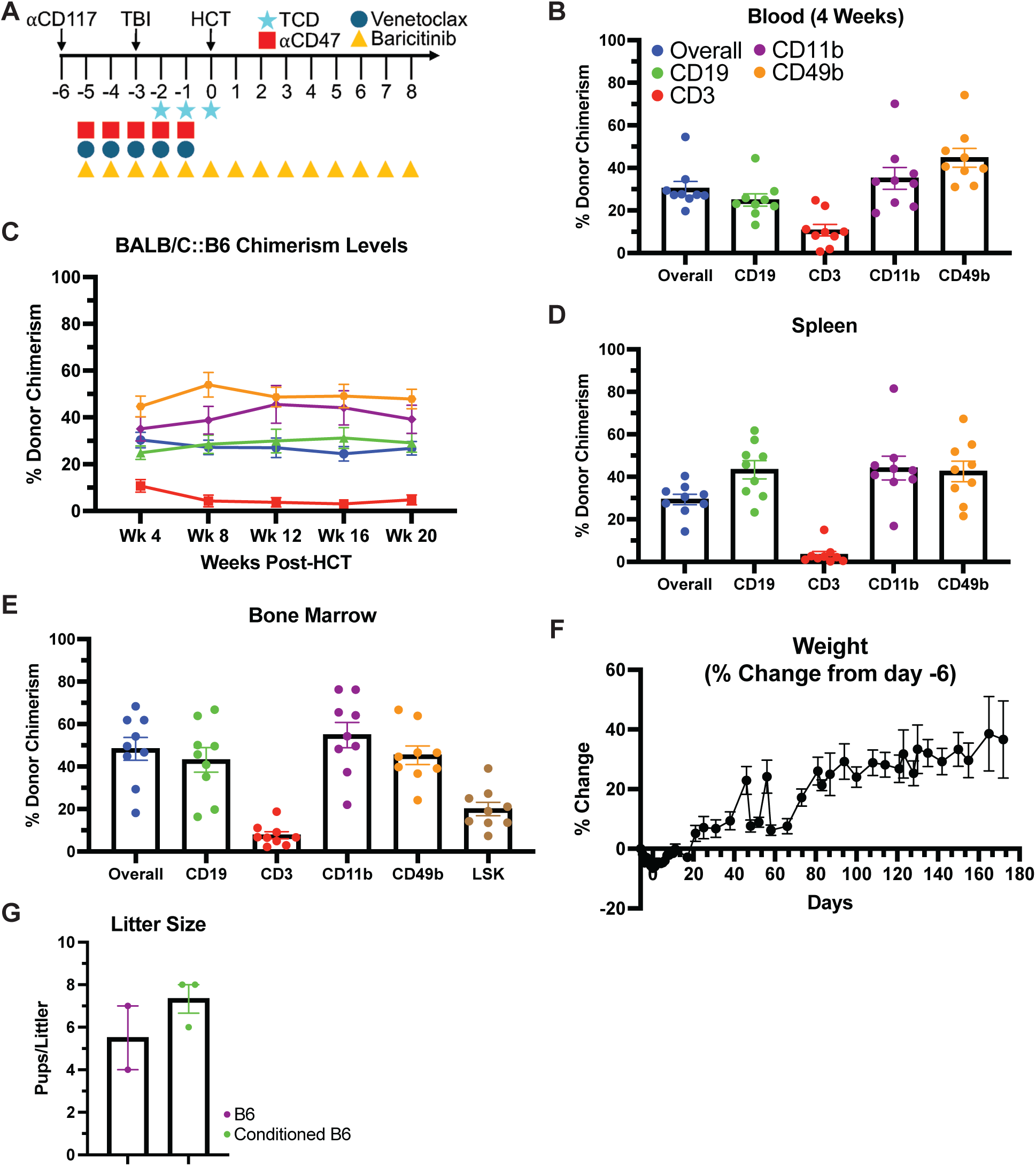
Durable mixed hematopoietic chimerism after non-myeloablative conditioning with 10cGy TBI. **(A)** Experimental conditioning and transplantation timeline. **(B)** Multilineage chimerism analysis of peripheral blood 4 weeks post-HCT. **(C)** Longitudinal multilineage chimerism analysis of peripheral blood through 20 weeks post-HCT. **(D)** Multilineage chimerism analysis of host spleen 22 or 25 weeks post-HCT. **(E)** Multilineage chimerism analysis, including Lin^-^Sca1^+^cKit^+^ (LSK) HSCs, of host bone marrow 22 or 25 weeks post-HCT. **(F)** Weight after HCT of BALB/c:B6 as a percentage of initial weight prior to conditioning start. (B-F) n = 9, 2 independent experiments. **(G)** Litter sizes of unconditioned female mice paired with conditioned males (purple; n = 2) and conditioned female mice paired with unconditioned males (green; n = 3). Data are represented with mean ± SEM. TBI = total body irradiation; HCT = hematopoietic cell transplant; TCD = T cell depletion. See also, **Supplementary Figure 4**.

While this conditioning regimen was well tolerated, mice developed peri-transplant anemia, corrected by a single whole blood transfusion. Despite the requirement for supportive intervention early on, conditioned mice showed normal average weight gain after conditioning alone, or conditioning and HCT. Importantly, both male and female mice remain fertile after conditioning, a significant finding and did not show indications of GVHD (**Figure 3G)**.

### Diabetes reversal with allogeneic islet and bone marrow transplantation after NMA conditioning

We investigated if combined allogeneic HCT and islet transplantation could reverse established diabetes in B6 RIPDTR mice after conditioning with αCD117, αCD47, TCD, venetoclax, baricitinib, and 10 cGy TBI (**Figure 4A**; see **Methods** (36)). After injection of diphtheria toxin (DT; red arrow), all B6 RIPDTR mice became severely diabetic, with glycemia >500 mg/dL (**Figure 4, E** and **F**). Prior to conditioning, diabetic mice were maintained on insulin (**Methods**) which was discontinued on day 0 after HCT and islet transplantation. We transplanted BALB/c WBM and ∼400 islets from BALB/c (donor-matched) or FVB (third-party) donors in the subcapsular renal space (**Figure 4A**). We observed multilineage mixed hematopoietic chimerism in peripheral blood of diabetic B6 RIPDTR by 4 weeks post-HCT (**Figure 4B**). Of five mice transplanted with BALB/c islets and followed for 20 weeks, all had durable chimerism (**Figure 4C**). Endpoint analysis of bone marrow revealed abundant donor chimerism, including LSK HSCs (**Figure 4D**). Thus, our NMA conditioning regimen incorporating only 10 cGy TBI achieved durable mixed hematopoietic chimerism and established allogeneic tolerance across full MHC mismatch in overtly diabetic mice.

**Figure 4:**
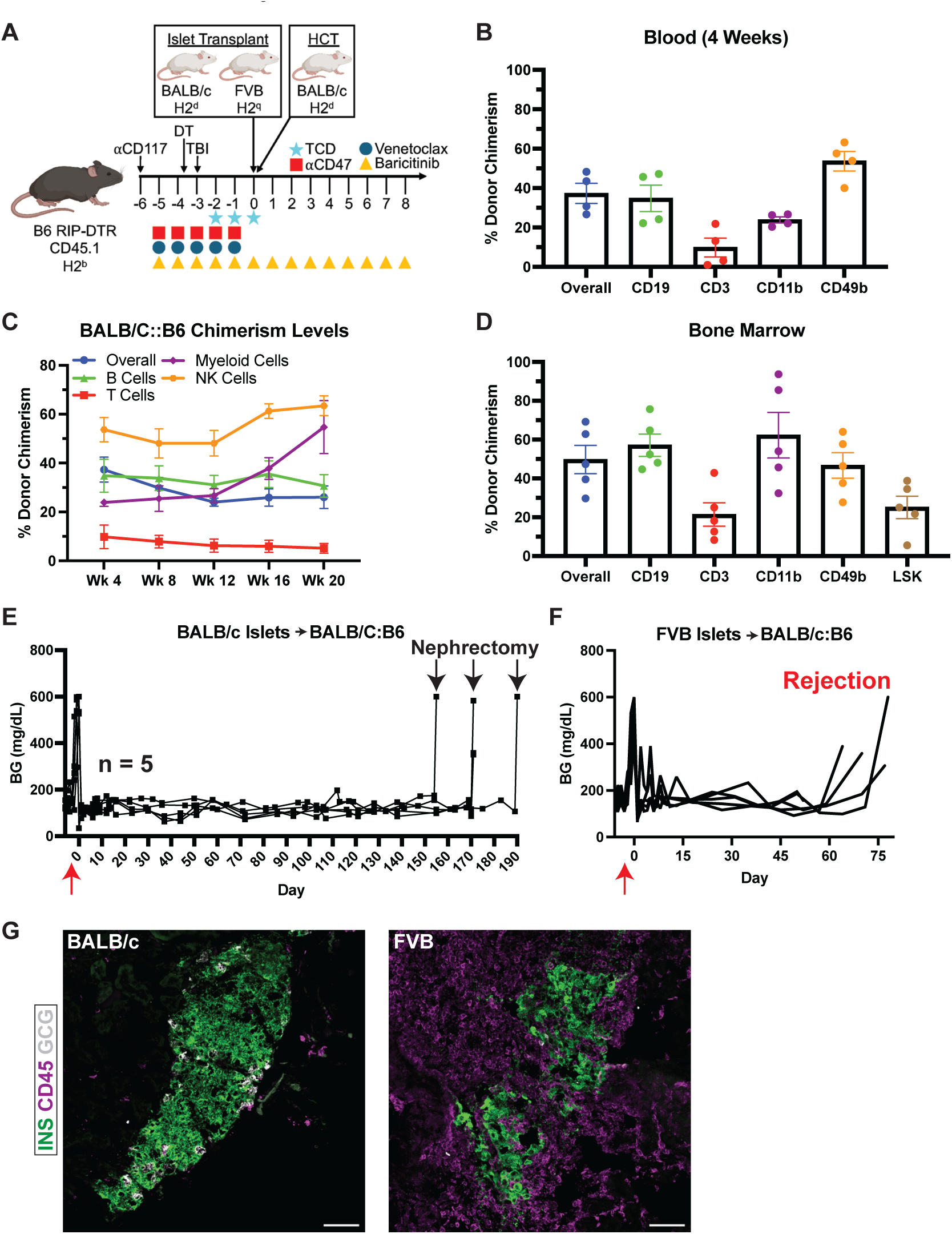
Allogeneic islet tolerance and diabetes reversal after simultaneous islet and bone marrow transplant in diabetic mice conditioned with 10 cGy TBI. **(A)** Experimental design and transplantation timeline. **(B)** Multilineage chimerism analysis of peripheral blood 4 weeks post-HCT. **(C)** Longitudinal multilineage chimerism analysis of peripheral blood through 20 weeks post-HCT. **(D)** Multilineage chimerism analysis, including Lin^-^Sca1^+^cKit^+^ (LSK) HSCs, of host bone marrow 24 or 27 weeks post-HCT. **(E)** Non-fasting blood glucose levels of diabetic B6 RIP-DTR mice that received simultaneous BALB/c bone marrow and islet transplants. (B-E) n = 5, 3 independent experiments. **(F)** Non-fasting blood glucose levels of diabetic B6 RIP-DTR mice transplanted with BALB/c bone marrow and third-party FVB islets (n = 5, 4 independent experiments). Red arrow indicates DT injection, black arrows indicate nephrectomy. (G) Representative maximum intensity projections of BALB/c (left) and FVB (right) islets transplanted under the kidney capsule of BALB/c:B6 mice stained for insulin (green), CD45 (magenta), and glucagon (white). Scale bar = 50 µm. Data are represented with mean ± SEM. TBI = total body irradiation; HCT = hematopoietic cell transplant; TCD = T cell depletion.

Diabetic B6 RIPDTR mice with durable chimerism (BALB/c:B6 RIPDTR) maintained normal blood glucose levels throughout the follow-up period (>150 days), and did not require supplemental insulin or additional immunosuppression (n=5/5; **Figure 4E**). We did not observe evidence of GVHD in chimeric BALB/c:B6 RIPDTR mice, and longitudinal body weight measures showed normal average weight gain after HCT.

Removal of the kidney harboring engrafted BALB/c islets (“Nephrectomy”; black arrows), resulted in rapid reversion to diabetes (n=5/5; **Figure 4E**). Chimeric BALB/c:B6 RIPDTR that received FVB islets had initial glycemic stabilization but spontaneously reverted to hyperglycemia by ∼70 days post-HCT, indicating rejection of these ‘third-party’ islets and re-establishment of immune competence (**Figure 4F**; n=5/5). Histology of the recovered islet graft at the experimental endpoint in BALB/c:B6 RIPDTR mice revealed intact BALB/c islets with little to no CD45^+^ immune cell infiltrate. By contrast, FVB islet grafts showed heavy immune cell infiltration and reduced islet cells (**Figure 4G**). In summary, we have developed a conditioning regimen with 10 cGy TBI that promotes durable mixed hematopoietic chimerism, allogeneic graft tolerance, and diabetes reversal.

### Central mechanisms of immune tolerance in mixed chimeric mice

Thymic dendritic cells (DCs) augment ‘negative selection’ of T cells in the thymus (37–40). In mixed chimerism, donor antigen presenting cells (APCs) in thymus, including thymic DCs, present both donor and self-antigens to developing thymocytes, promoting autologous and allogeneic tolerance(41). To assess tolerance mechanisms in BALB/c:B6 chimeras, we characterized the major CD11c^+^ dendritic cell subsets: B220^+^PDCA1^+^ plasmacytoid DCs (pDCs), conventional CD8^+^SIRPα^-^ cDC1 cells, and CD8^-^SIRPα^+^ cDC2 cells in the thymus and spleen of chimeric BALB/c:B6 mice compared to controls (37–40,42). We confirmed the presence of CD45.2^+^ donor-derived DCs in both the thymus (**Figure 5A**) and spleen (**Figure 6A**) of mixed chimeric mice, indicative of antigen presentation by donor DCs in these tissues. Compared to both unconditioned and conditioning-only control B6 mice, we detected thymic pDC and cDC2 DCs at similar frequencies in BALB/c:B6 mice (**Figure 5B**). Interestingly, the frequency of cDC1s was significantly increased in conditioned controls relative to BALB/c:B6 chimeric mice with or without an islet graft (**Figure 5B**). This difference in cDC1 frequency highlights potential shifts in immune regulation, warranting further exploration of pathways like the PD-1/PD-L1 signaling axis, which is critical for maintaining central and peripheral tolerance (43,44).

**Figure 5:**
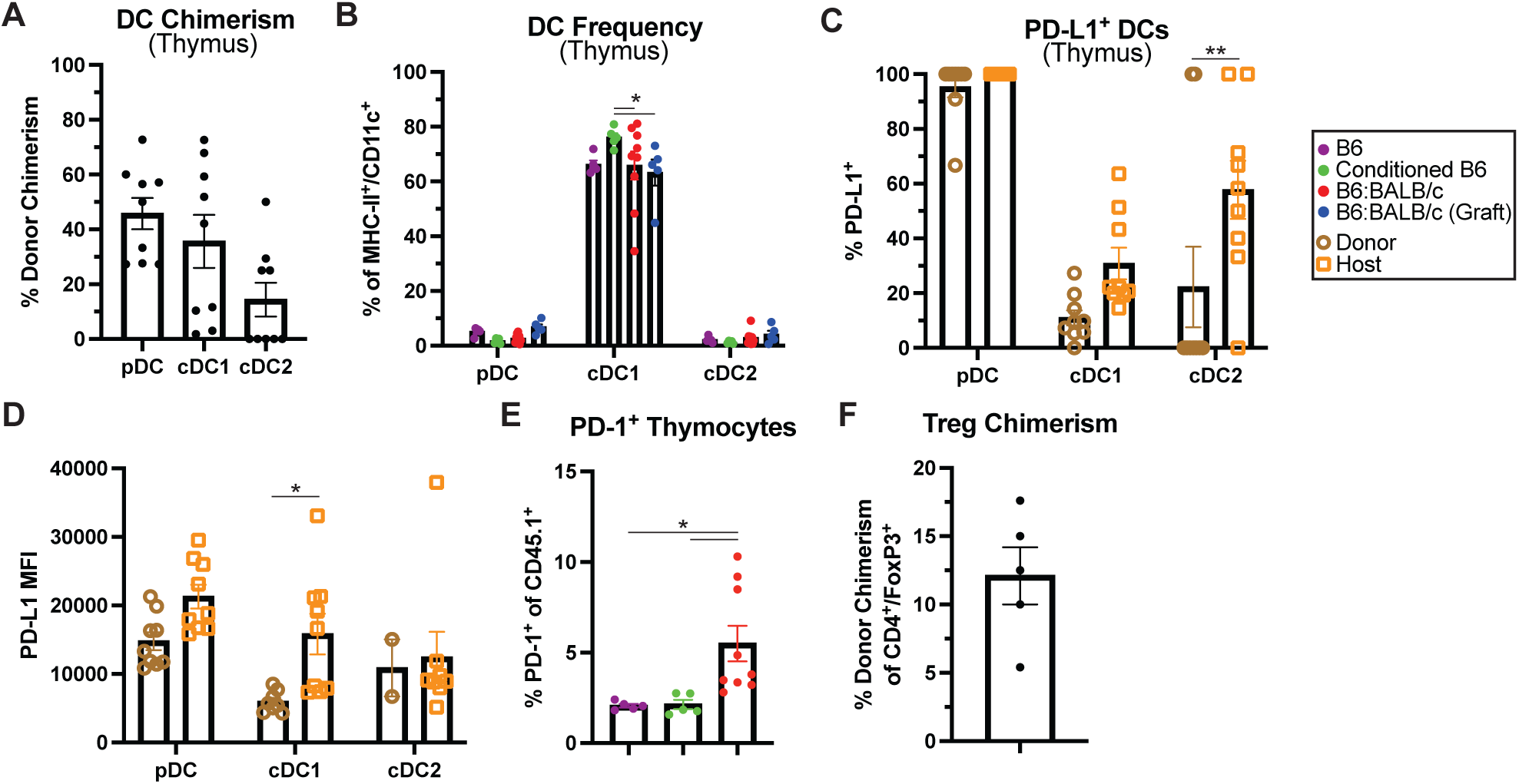
Central tolerance mechanisms in mixed chimeric mice. **(A)** Chimerism analysis of CD11c^+^MHC-II^+^ dendritic cell subsets in the thymus 22 or 25 weeks post-HCT. pDCs = B220^+^PDCA1^+^, cDC1 = B220^−^SIRP⍺^−^ CD8^+^, cDC2 = B220^−^SIRP⍺^+^CD8^−^. **(B)** Thymic DC subset frequency in WT B6, conditioned B6, B6/BALB/c mixed chimeras, and BALB/c:B6 mixed chimeras that received an islet graft (n = 5-9). Proportion **(C)** and MFI **(D)** of PD-L1 expression in thymic host- and donor-derived DCs in BALB/c:B6 mice (*n* = 9). **(E)** Proportion of CD45.1^+^/PD-1^+^ thymocytes in BALB/c:B6, B6 and conditioned B6 control mice (n = 5-9). **(F)** Donor chimerism in CD4^+^/FoxP3^+^ thymic Tregs (n = 5). Data presented as mean ± SEM. *P < 0.05, **P < 0.01, ***P < 0.001, ****P < 0.0001. See also **Supplementary Figure 5**.

**Figure 6:**
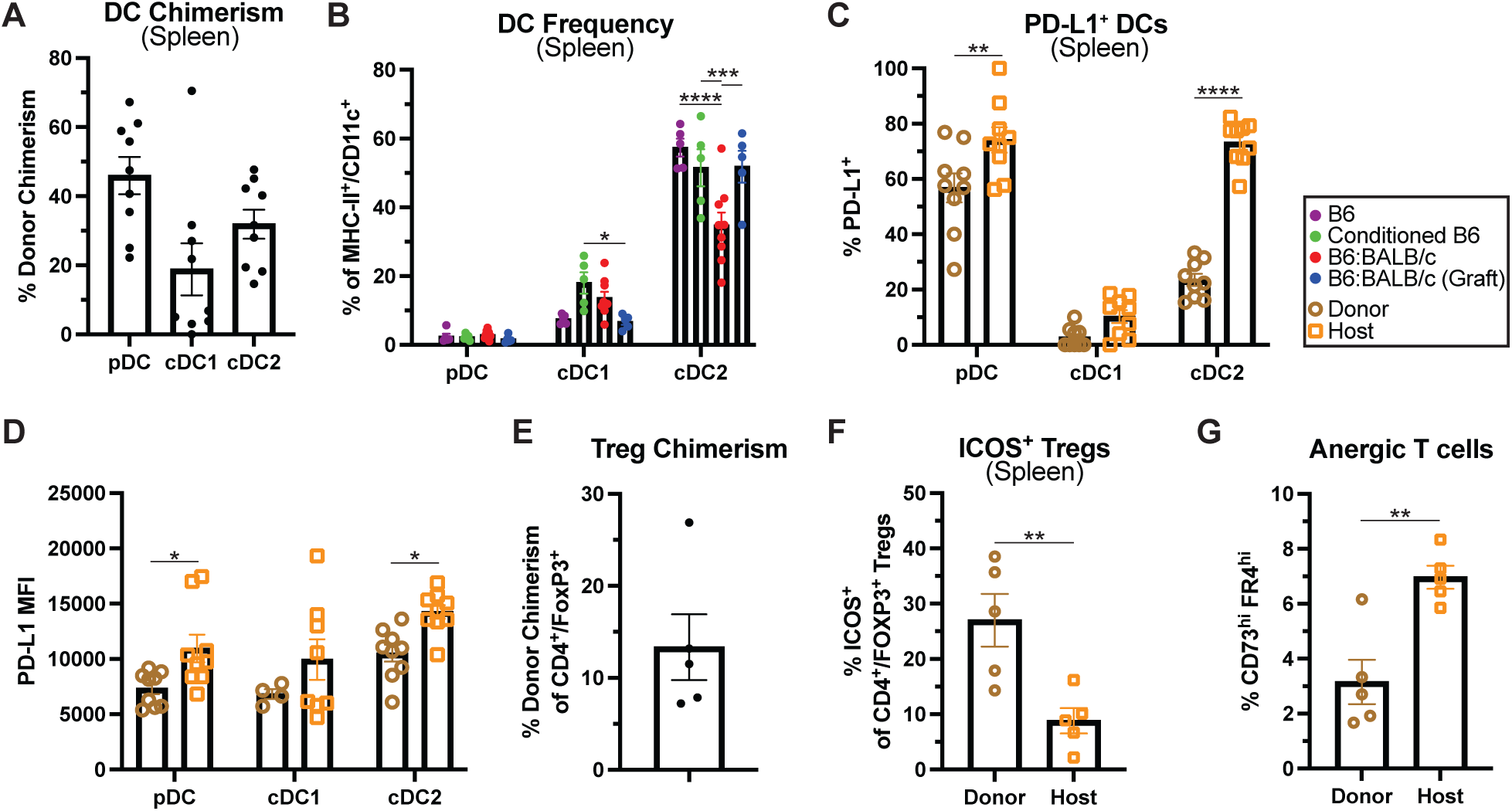
Peripheral tolerance mechanisms in mixed chimeric mice. **(A)** Chimerism analysis of CD11c^+^MHC-II^+^ dendritic cell subsets in the spleen 22 or 25 weeks post-HCT. pDCs = B220^+^PDCA1^+^, cDC1 = B220^−^SIRP⍺^−^CD8^+^, cDC2 = B220^−^SIRP⍺^+^CD8^−^. **(B)** Splenic DC subset frequency in WT B6, conditioned B6, B6/BALB/c mixed chimeras, and BALB/c:B6 mixed chimeras that received an islet graft (n = 5-9). Proportion **(C)** and MFI **(D)** of PD-L1 in splenic host and donor DCs in BALB/c:B6 mice (*n* = 9). **(E)** Donor chimerism in CD4^+^/FoxP3^+^ splenic Tregs (n = 5). **(F)** Proportion of ICOS^+^/CD4^+^/FoxP3^+^ splenic Tregs (n = 5). **(G)** Frequency of CD73^hi^ FR4^hi^ anergic cells among host and donor CD4^+^ FOXP3^-^ T_con_ cells of BALB/c:B6 spleen. Data presented as mean ± SEM. *P < 0.05, **P < 0.01, ***P < 0.001, ****P < 0.0001

In the thymus, PD-1 is upregulated in thymocytes undergoing negative selection and primarily localized to the medulla, while PD-L1 is expressed in both the cortex and medulla by thymic epithelial and dendritic cells (44–46). We observed an increase in the frequency and levels of PD-L1 expression in host-derived cDC1 and cDC2 DCs of mixed chimeric mice (**Figure 5, C** and **D**). We also observed significantly increased overall frequency of PD-1^+^ thymocytes in chimeric mice, compared to controls (**Figure 5E**). To further evaluate the selection of donor and host-derived thymocytes, we analyzed T cells with the Vβ11 domain, which are deleted by genome-encoded superantigen presentation in BALB/c but not B6 mice (47). In BALB/c:B6 mice, ∼70% of Vβ11^+^ T cells were depleted, compared to B6 controls, indicating education of donor T cells to host antigens (**Supplementary Figure 5A**). By contrast, Vβ8.1^+^ T cells, which are not under selective pressure in BALB/c or B6 strains, were not depleted (**Supplementary Figure 5B**). Negative selection of thymocytes is often accompanied by the presence of thymic regulatory T cells (tTregs), and we detected donor-derived CD4^+^/FoxP3^+^ tTregs in the thymus of chimeric BALB/c:B6 mice (**Figure 5F**). Together, these results support mechanisms of central tolerance induction in BALB/c:B6 mixed chimeras, where donor DCs in the host thymus could mediate negative selection and augment the generation of host and donor-derived tTregs.

### Peripheral mechanisms of immune tolerance in mixed chimeric mice

Peripheral tolerance mechanisms are crucial for suppressing allo- and autoreactive T cells that evade thymic selection and for tolerizing residual tissue-resident host T cells that survive conditioning. As with thymic DCs, we evaluated both the frequency of splenic DC subsets and their PD-L1 expression (**Figure 6, B-D**). Similarly, we observed a significant increase in the frequency and level of PD-L1 expression in host-derived splenic pDCs and cDC2s (**Figure 6, C** and **D**). These findings indicate a role for PD-L1 expression in splenic DCs in promoting peripheral tolerance.

Peripheral tolerance in transplantation is also associated with Treg activity (48). Consistent with our observations in the thymus, we observed donor-derived splenic CD4^+^/FoxP3^+^ Tregs in mixed chimeric mice (**Figure 6E**). To determine whether Tregs present in mixed chimeric mice were functional, we analyzed inducible costimulator (ICOS), whose expression is correlated with increased IL-10 secretion and suppression potential, in donor- and host-derived splenic Tregs (49). While both host- and donor-derived ICOS^+^ Tregs are present, donor-derived ICOS^+^ Tregs are significantly more abundant than host (**Figure 6F**). Donor-derived Tregs can suppress alloreactive host-type T cells (50). Supporting this possibility, we observed that the proportion of host-derived anergic CD4^+^/CD73^hi^/FR4^hi^ cells was significantly higher (**Figure 6G**). In summary, durable mixed hematopoietic chimerism after conditioning with our low-dose radiation regimen likely reflects both central and peripheral mechanisms that establish and maintain tolerance to donor HSCs and islets and prevent GVHD.

## Discussion

Mixed hematopoietic chimerism can induce allogeneic, donor-specific tolerance across MHC mismatched barriers. Indeed, in patients suffering renal failure who received donor-matched HCT and kidney allotransplants, this approach has achieved decades-long renal replacement without systemic immunosuppression (8,10,51,52). Despite demonstrated proof-of-concept in humans, broader use of mixed hematopoietic chimerism for tolerance induction is limited by the toxicities associated with conventional HCT transplant protocols that use high doses of radiation or DNA-damaging chemotherapy (41,53). Pre-transplant conditioning is necessary to create niche space in the host bone marrow donor HSCs to engraft and to prevent immune rejection of transplanted cells (14). Thus, we developed a NMA condition regimen that supports durable multi-lineage mixed hematopoietic chimerism after nearly complete elimination of conditioning irradiation. Transplantation of donor-matched islets in diabetic, immune competent mice, resulted in durable diabetes reversal without chronic immunosuppression or complications like GVHD. This represents a significant advance from our prior work (23).

Chemotherapeutics and radiation doses currently used for HCT conditioning are associated with myriad morbidities, including impaired endocrine function, fertility loss, and increased risk for secondary malignancy (54). Here, we demonstrate that the non-genotoxic conditioning agents incorporated in our conditioning regimen are well tolerated, do not affect pancreatic islet function, and maintain other measures of functional status, like fertility. A single dose of TBI <10 cGy is operationally defined as ‘low dose’ and is not associated with increased carcinogenic risk(55); thus, we sought to reduce conditioning TBI to ≤10 cGy, with the aspiration to broaden adoption of mixed chimerism for organ replacement. To achieve donor cell engraftment with reduced irradiation, we systematically assessed conditioning reagents currently in clinical use or involved in clinical trials. Baricitinib (56), venetoclax (57), αCD4 monoclonal antibody (ibalizumab) (58), and NMA radiation are all currently approved for the treatment of multiple non-malignant or malignant disease. Human αCD117 Ab (JSP191) is in multi-center trials (ClinicalTrials.gov: NCT02963064, NCT04429191, NCT04784052) and has shown a good safety profile and promising results. Likewise, numerous αCD47 monoclonal antibodies are currently under clinical evaluation (59). While the development and clinical evaluation of αCD8 monoclonal antibodies is limited, transient T cell depletion can also be achieved with anti-thymocyte globulin or other conditioning agents currently in clinical use.

GVHD is a potentially life-threatening complication of HCT and an impediment to expanding HCT for non-malignant disorders (60). Conditioning intensity and HSC sourcing and composition are major components of GVHD risk (61–63). While enriched HSPC populations, depleted of mature effector cells, can be used to mitigate GVHD risks associated with WBM preparations, enriched HSPCs do not engraft as well as WBM (64,65). We achieved similar chimerism outcomes with our conditioning regimen and either WBM or lineage-depleted HSPC transplantation, and observed no signs of GVHD like hair loss, rashes, postural changes, or weight loss following either. On the contrary, longitudinal body weight measures showed excellent weight gain from HCT to experimental endpoint in all comparison groups. Baricitinib use to prevent and treat GVHD has been evaluated in animal models, and is currently being evaluated for clinical treatment of GVHD (66,67). Thus, in addition to being a key element of our NMA conditioning regimen, baricitinib might both suppress peri-transplant acute GVHD, and prevent chronic GVHD.

To assess mechanisms of allotolerance in B6 mice with mixed chimerism, we characterized cellular and molecular features of central and peripheral tolerance. We observed durable mixed chimerism in all immune cell lineages and compartments evaluated, including the bone marrow, spleen, thymus and circulating blood. Furthermore, the data support a role for thymic-mediated central tolerance, including evidence of donor APC chimerism and elimination of host-reactive Vβ11^+^ T cells, in promoting allogeneic tolerance in these settings. Lack of islet immune cell infiltration in mixed chimeric B6 RIPDTR mice after HCT and islet transplantation suggests that donor alloantigen tolerance was achieved using mixed chimerism. Maintenance of immune competence after NMA conditioning and islet transplantation to achieve diabetes reversal is an important goal for clinical translation. We observed robust rejection of third-party FVB islet grafts by mixed chimeric BALB/c:B6 mice, indicating reconstituted immune function; however, additional provocative challenges are required to assess the immune status of BALB/c:B6 mice further.

In summary, we demonstrate the use of a clinically portable NMA conditioning regimen and islet transplantation protocol to establish mixed hematopoietic chimerism and allogeneic tolerance across full MHC barriers. Features of this strategy, including a unique combination of conditioning agents with synergistic mechanisms that minimize XRT, provide a significant conceptual advance for islet allotolerance, and could promote its clinical adoption in diabetes. Importantly, these findings could be expanded and applied to induce tolerance of other solid organ transplants, eliminating the need for chronic immune suppression after transplant. Expanding on these findings, further work will evaluate whether this strategy could be adapted for use in reversing autoimmune diabetes, or other autoimmune disorders, and to induce tolerance to replacement islet cells derived from renewable sources, including multipotent stem cells.

## Methods

### Sex as a biological variable

Our study examined male and female animals, and similar findings are reported for both sexes.

### Animals

Female and male B6 CD45.1 (Stock #: 002014), BALB/c (Stock #: 000651), and FVB (Stock #: 001800) mice were purchased from The Jackson Laboratory (Bar Harbor, ME). B6 *RIP-DTR* mice were generated and maintained by our group and used at 10-20 weeks of age(36). This strain expresses the *Ins2-HBEGF* (RIP-DTR) transgene and the mutant *Ptprca* (CD45.1) allele on the B6 mouse background. The RIP-DTR transgene allows for rapid induction of diabetes by β cell-specific ablation and 100% penetrance with a single i.p. injection of diphtheria toxin in males and females. Healthy euglycemic littermates of the same sex were randomly assigned to experimental groups and were not involved in any prior procedures. All animals were fed standard chow and water *ad libitum* and housed in non-specific-pathogen-free conditions at the Stanford School of Medicine. Animal experiments were approved by the Stanford Administrative Panel on Laboratory Animal Care (IACUC).

### Conditioning, reagents, and equipment

A graphical timeline of conditioning is shown in Figure 3A. Mice were given 500μg diphenhydramine HCl intraperitoneally (i.p.) approximately 10-15 minutes prior to αCD117. 500μg αCD117 (95934, BioLegend) was injected retro-orbitally into mice under isoflurane anesthesia on day −6 prior to HCT. Mice were irradiated on day −3 TBI. 300μg each of αCD4 and αCD8 was administered i.p. on days −2, −1, and 0. The selective JAK1/2 inhibitor, baricitinib, was dissolved in 100% DMSO at 20 mg/mL and stored at –20°C in 100 μL aliquots. DMSO stocks were thawed and diluted 1:10 in PBS immediately prior to use, and 200 μL/mouse was injected subcutaneously (s.c.) (400 μg). Baricitinib was dosed days −5 to +3 or +8, according to timelines. αCD47 was administered days −6 to −2 (Figure 2, S3) or days −5 to −1 (Figure 3, 4, S4) at the stated concentrations. The selective Bcl-2 inhibitor, venetoclax, was dissolved in 100% DMSO at 20 mg/mL and stored at –20°C in 50 μL aliquots. Immediately prior to use, DMSO stocks were thawed and diluted in 400 μL PEG-300, 50 μL Tween 80, 500 μL ddH_2_O; solvents added in this order and mixed thoroughly in-between. 200 μL/mouse (200 μg) venetoclax was injected i.p. Venetoclax was administered days −5 to −1. Animal irradiation (XRT) was performed in a Kimtron Polaris IC-250 Biological Irradiator (Oxford, CT) with a 225 kV X-ray tube filtered by 0.5mm Cu source set at 225kV, 13.3mA. Mice were divided in irradiation pie cages from Braintree Scientific (Braintree, MA) when irradiated. Dosimetry calibration for our setup was performed using published methods on radiochromic film dosimetry (Ma et al., 2001).

### Bone marrow isolation, enrichment, and transplant

Donor BALB/c mice (6-7 weeks old) were euthanized, and femurs, tibias, and vertebral bodies were collected. Bones were crushed via mortar and pestle in PBS with 2% FBS, 10mM HEPES, and 2mM EDTA to recover WBM. WBM was filtered through a 70-µm cell strainer, and RBCs were lysed in RBC Lysis Buffer (BioLegend). WBM cells were stained with Trypan Blue (StemCell Technologies; Vancouver, Canada) and counted with a Countess 3 Automated cell counter (ThermoFisher Scientific; Waltham, MA). Lineage-negative (Lin-) bone marrow cells were enriched by magnetic column separation using a Lineage Cell Depletion cocktail (Miltenyi Biotec; Bergisch Gladbach, Germany) as per manufacturer’s instructions. 1.5E6 Lin-cKit+ cells or 30E6 WBM cells resuspended in 100 µl of PBS were injected retro-orbitally. Lin-cKit+ HSPC preparation composition has been previously described(23).

### Islet isolation and transplantation

Islet isolation and transplantation was performed as previously described with minor modifications(68,69). Briefly, pancreases are perfused with 100-125μg/mL Liberase TL (Roche Diagnostics; Indianapolis, IN) through the common bile duct and digested in a 37°C water bath for 18-22 minutes. After washing with Hank’s Buffered Saline (HBS; Caisson Labs; Smithfield, UT), the crude digest is purified over a discontinuous density gradient, washed once more with HBS, and cultured overnight in 5.5 mM glucose RPMI 1640 (Corning; Corning, NY) supplemented with 10% FBS, 10mM HEPES, and 1% penicillin-streptomycin solution. Recipient mice were anesthetized with a ketamine/xylazine mix and given subcutaneous analgesics. 200-400 islets were injected under the kidney capsule of recipient mice with a micro-capillary, as described(70). The nephrectomy procedure involved the same anesthetic regimen as islet transplantation and renal vessels were first tied to prevent hemorrhage before the kidney containing islet graft was removed.

### Histology

Islet graft-bearing kidneys, pancreases, and intestines were fixed in 4% paraformaldehyde overnight, incubated overnight in 30% sucrose, embedded in optimal cutting temperature compound, and frozen on dry ice. 6-10μm sections were made on a Leica CM3050 S (Leica Biosystems; Buffalo Grove, IL). Immunofluorescent staining was performed using standard methods. Briefly, sections were blocked for 1hr then incubated with primary antibodies overnight at 4°C. Sections were washed for 5 minutes x 3 before incubation with secondary antibodies for 2 hours at room temperature or overnight at 4°C and washed 3 x 5 minutes again. Slide covers were set with Hard-set Mounting Medium with DAPI (Santa Cruz Biotechnology; Dallas TX). Slides were imaged on Zeiss AxioM1 or Leica SP2 confocal microscopes. Post-processing and color channel merging was performed in Fiji (http://fiji.sc/) (71). Primary and secondary antibodies and dilutions are documented (Key Resources Table).

### Peripheral blood, spleen, thymus, and BM preparation for flow cytometry

100 mL of whole blood was collected via the tail vain into EDTA coated tubes. Spleens and thymuses were directly mashed through a 70mm cell strainer. BM cells were isolated as above. Samples underwent RBC lysis in RBC Lysis Buffer (BioLegend) for 5-10 min at 4°C before down-stream staining for analysis.

### Flow cytometry analysis

Gating strategies were previously described(23). For analysis of mixed chimerism, cells were first stained with LIVE/DEAD Fixable Near-IR Dead Cell Stain Kit (ThermoFisher Scientific) and blocked with TruStain FcX anti-mouse (BioLegend) for 10 min on ice in Cell Stain Buffer (BioLegend). Antibodies used for staining were from BioLegend. Extracellular markers were stained with antibodies listed (Key Resources Table) at manufacturer’s recommended dilutions. Staining of intracellular markers was conducted with BioLegend True-Nuclear Transcription Factor Buffer Set as per manufacturer’s instructions. Cells were analyzed with a 5L Aurora (Cytek Biosciences; Fremont, CA). Data were analyzed using FlowJo (10.9).

### Statistics

Statistical details of all experiments can be found in the figure legends and results section, including value of n. All data are presented as means ± SEM, where n represents number of animals. Animals were randomly assigned to experimental groups, and all samples represent biological replicates. Statistical analysis was performed using Prism 10 (GraphPad, San Diego, CA). Differences between the means of two groups were tested using unpaired two-tailed Student’s t-test with Welch’s correction. Differences between the means of three or more groups were tested by 2way ANOVA using Tukey’s Multiple Comparisons test with Geisser-Greenhouse correction. Some data were excluded by Prism 10’s outlier function. Sample size estimates were not used. A p value of 0.05 or less was considered statistically significant. *P < 0.05, **P < 0.01, ***P < 0.001, ****P < 0.0001.

### Study approval

Animal experiments were approved by the Stanford Administrative Panel on Laboratory Animal Care, in line with ARRIVE guidelines.

## Data Availability

All data generated or analyzed during this study are included in this published article (and its supplementary information files) or are available from the corresponding author on reasonable request. The supporting data values file includes values underlying graphed data and reported means presented in both the main text and supplemental material.

## Author Contributions

S.A.R. designed and performed experiments, data collection, data analysis, wrote the manuscript and was involved in funding acquisition. P.B. advised on experimental design and assisted in performing experiments, data collection, data analysis and visualization, edited the manuscript, and was involved in funding acquisition. D.M.B performed experiments, data collection, data analysis and visualization. X.G., R.R., N.N, and M.N. performed experiments and data collection. J.A.S., and S.P. provided guidance and feedback on experimental design, results, provided reagents, reviewed and edited the manuscript, and were involved in funding acquisition. S.K.K. designed experiments, wrote the manuscript, supervised the project, acquired funding, and is the guarantor of this work.

## Acknowledgments

We thank the members of the Kim group, especially Drs. Y. Hang, R. Whitener, and S. Park, and Dr. E. Meyer (Stanford) for advice and encouragement. We especially thank Ms. L. Nichols, director of the Stanford Shared FACS Facility, where flow cytometry analysis was performed. CBC testing was done by the Stanford Animal Diagnostic Laboratory. Imaging in this study was performed at the Stanford University Cell Science Imaging Facility (RRID: SCR_017787). We thank the Stanford Diabetes Research Center (SDRC) Islet Core, especially Ms. X. Gu and Dr. J. Wang, for performing islet isolation, islet transplantation, and nephrectomies. S.A.R. is supported by the Institute for Immunity, Transplantation and Infection – Stanford Autoimmunity & Allergy Supergroup and the NIH (LAUNCH 1TL1DK139565-0, F32DK141209). P.B. is a student in the Medical Scientist Training Program (MSTP) and PhD Program in Immunology at Stanford. P.B. is supported by the NIH (T32 GM736543) and the Stanford Interdisciplinary Graduate Fellowship through Bio-X (Morgridge Family Fellow). D.M.B. is supported by the VPUE Research Fellowship at Stanford. M.N. is supported by a Breakthrough T1D Postdoctoral Fellowship. Work in the Kim group is supported by the Breakthrough T1D Northern California Center of Excellence (S.K.K and J.A.S.), NIH awards (R01 DK107507; R01 DK108817; U01 DK123743; P30 DK116074 to S.K.K), the Reid Family, H.L. Snyder Foundation and Elser Trust, and the Stanford Diabetes Research Center (SDRC). Work here was also supported by the Islet Research Core in the SDRC. Dr. Kim is the KM Mulberry Endowed Professor at Stanford University School of Medicine.

