## Supplementary Material for "Improved conditioning for hematopoietic chimerism induces islet tolerance to cure diabetes"

**Conflict of Interest**

S.A.R is a consultant and stockholder of Tolerance Bio, Inc. J.A.S. is a co-founder, stockholder, and board member of Jasper Therapeutics, Inc.

### Supplementary Materials

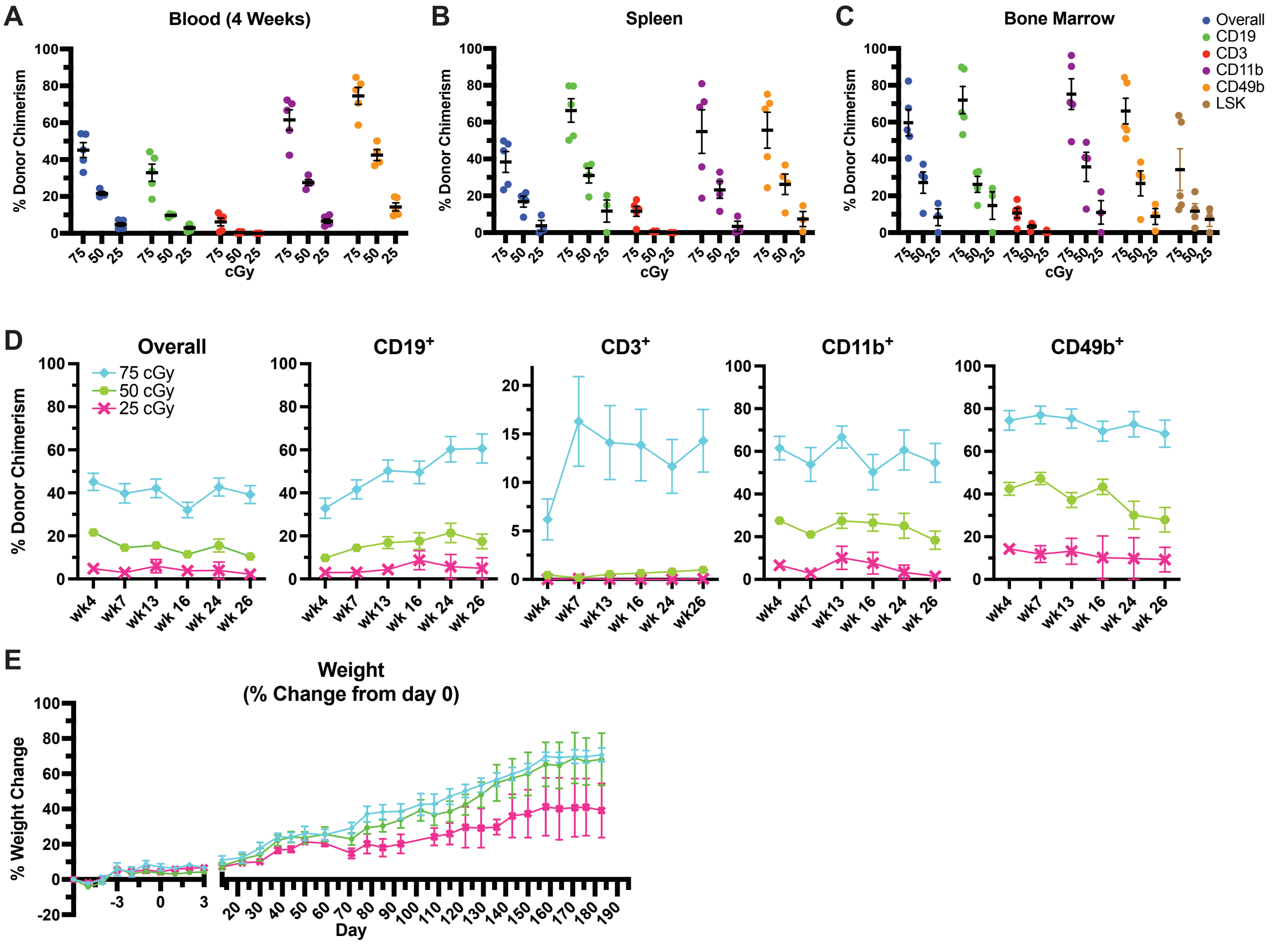


**Supplementary Figure 1: Lack of durable mixed hematopoietic chimerism after conditioning with baricitinib and ≤50 cGy TBI. (A)** Multilineage chimerism analysis of peripheral blood 4 weeks post-HCT in non-diabetic B6 mice conditioned as outlined in Fig 1B with 75, 50, or 25 cGy TBI (n = 5, 4, and 5 for 75, 50, and 25 cGy groups, respectively). **(B)** Multilineage chimerism analysis of host spleen 26 weeks post-HCT. **(C)** Multilineage chimerism analysis, including Lin$^{-}$Sca1$^{+}$cKit$^{+}$ (LSK) HSCs, of host bone marrow 26 weeks post-HCT. **(D)** Longitudinal multilineage chimerism analysis of peripheral blood through 26 weeks post-HCT. **(E)** Weight after conditioning and HCT as a percentage of starting weight. (C-E) n = 5, 4, and 3 for 75, 50, and 25 cGy groups, respectively. Data presented as mean ± SEM.

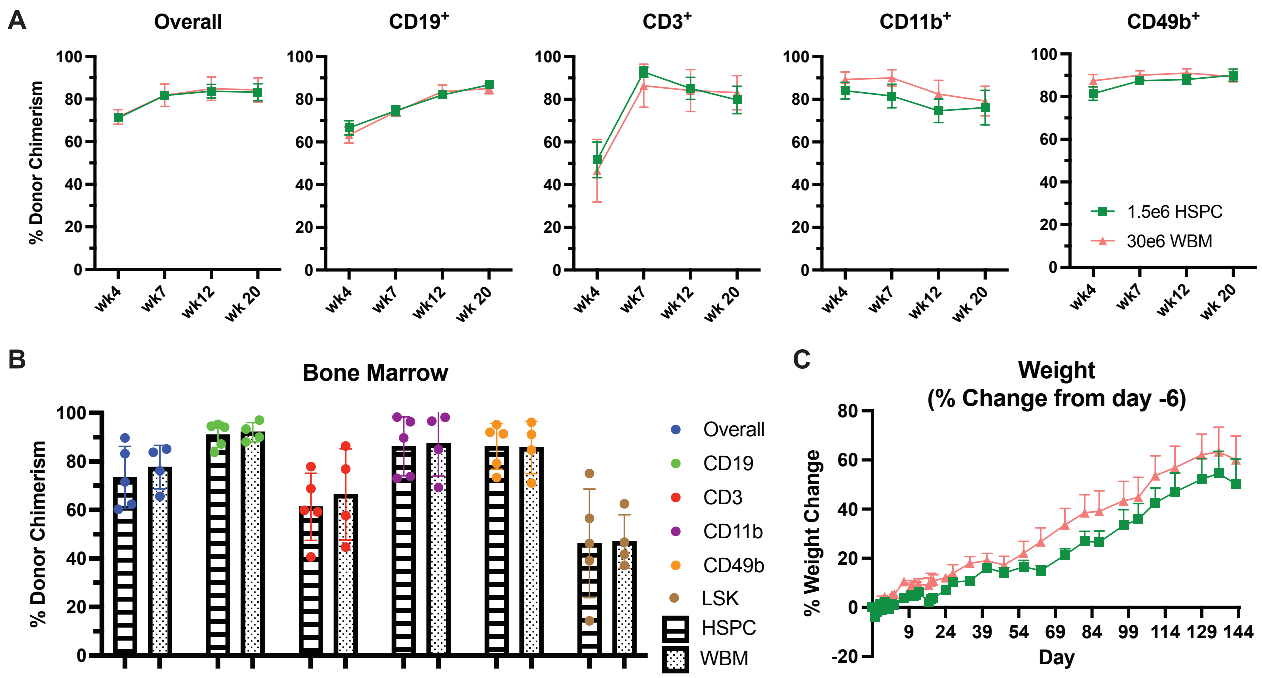


**Supplementary Figure 2: HCT source material does not affect chimerism levels in conditioned B6. (A)** Longitudinal multilineage chimerism analysis of peripheral blood through 20 weeks post-HCT. B6 mice were conditioned as outlined in Fig 1B and 200 cGy TBI and transplanted with 1.5e6 enriched HSPCs or 30e6 WBM cells. **(B)** Multilineage chimerism analysis, including Lin$^-$Sca1$^+$cKit$^+$ (LSK) HSCs, of host bone marrow 21 weeks post-HCT. (A-B) n = 5 and 4 for HSPC and WBM groups, respectively. Data presented as mean ± SEM.  HSPC = hematopoietic stem and progenitor cells; WBM = whole bone marrow.

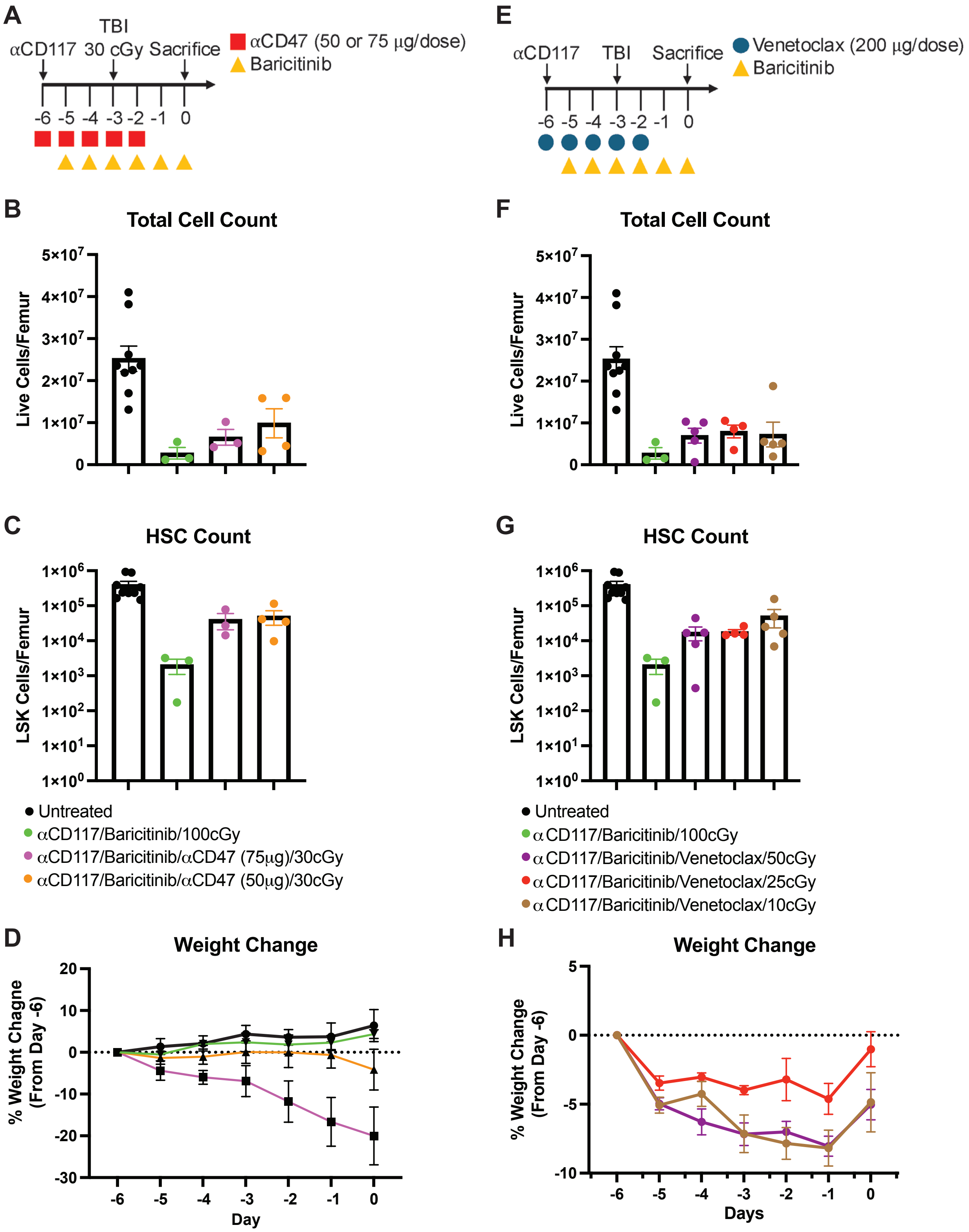


**Supplementary Figure 3: Evaluation of bone marrow clearance after conditioning with aCD47 antibody and Bcl2 inhibitor. (A)** Experimental conditioning timeline evaluating 5 daily doses of 50 or 75 µg/dose of aCD47 antibody. **(B)** Total live cell and **(C)** LSK cell counts per femur in unconditioned B6 CD45.1 mice and mice conditioned with αCD117/BTB/100 cGY TBI, αCD117/BTB/αCD47 (75 µg/dose)/30 cGy TBI, or αCD117/BTB/αCD47 (50 µg/dose)/30 cGy TBI. **(D)** Body weight change throughout conditioning period of as a percentage of initial weight at start of conditioning. **(E)** Experimental conditioning timeline evaluating 5 daily doses of Venetoclax and 50, 25, or 10 cGy TBI. **(F)** Total live cell and **(G)** LSK cell counts per femur in unconditioned B6 CD45.1 mice and mice conditioned with αCD117/BTB/100 cGY TBI, αCD117/BTB/venetoclax/50 cGy TBI, αCD117/BTB/venetoclax/25 cGy TBI, or αCD117/BTB/venetoclax/10 cGy TBI. **(H)** Body weight change throughout conditioning period of as a percentage of initial weight at start of conditioning. Data presented as mean ± SEM.

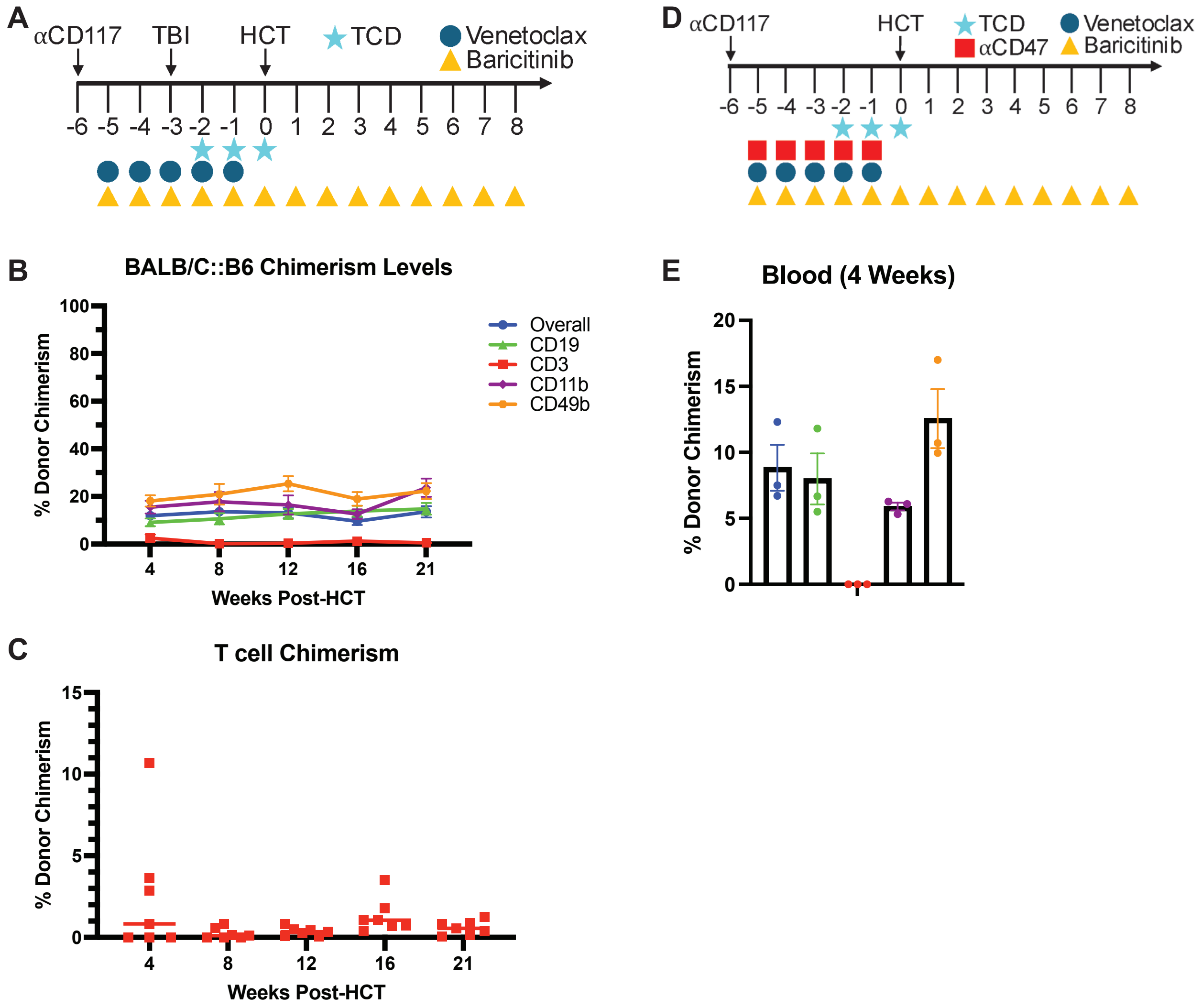


**Supplementary Figure 4: Non-myeloablative conditioning requires aCD47 and 10 cGy TBI durable allogeneic donor T cell chimerism.** **(A)** Conditioning timeline for B6 CD45.1 mice conditioned with αCD117/BTB/venetoclax/10 cGy TBI and no αCD47. **(B)** Longitudinal multilineage chimerism analysis of peripheral blood through 21 weeks post-HCT. **(C)** Individual donor T cell proportion data points over time. (B-C) n = 7. **(D)** Conditioning timeline for B6 CD45.1 mice conditioned with αCD117/BTB/αCD47/venetoclax and *no TBI*. **(E)** Multilineage chimerism analysis of peripheral blood 4 weeks post-HCT. TBI = total body irradiation; HCT = hematopoietic cell transplant. Data presented as mean ± SEM.

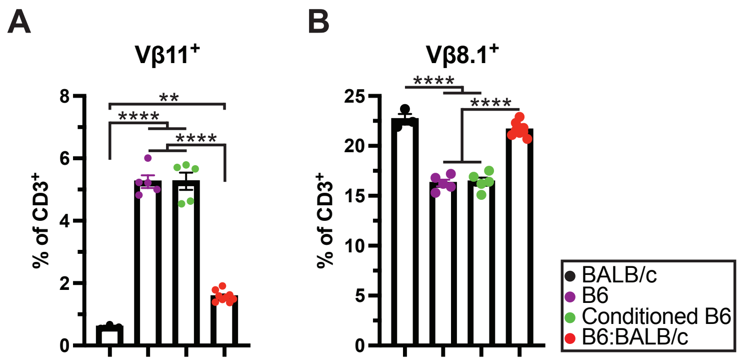


**Figure S5: Deletion of donor reactive thymocytes in BALB/c:B6 mice: (A)** Vβ11$^{+}$ and **(B)** Vβ8.1$^{+}$ splenic T cells in BALB/c, B6, conditioned B6 controls, and BALB/c:B6 mice (n = 3-9). Data presented as mean ± SEM. **P < 0.01, ***P < 0.001, ****P < 0.0001.

**Supplementary Table 1**

| REAGENT or RESOURCE | SOURCE | IDENTIFIER |
|---|---|---|
| **Antibodies** | | |
| Goat α-Guinea Pig CF594 | MilliporeSigma | Cat#: SAB4600103 |
| Goat α-Guinea Pig CF488A | MilliporeSigma | Cat#: SAB4600040 |
| α-Glucagon | ThermoFisher Scientific | RRID: AB_2804644<br>Cat#: PA5-88091 |
| α-Insulin | Dako | RRID: AB_10013624<br>Cat#: A0564 |
| α-CD45 | Biolegend | RRID: AB_312966<br>Cat#: 103102 |
| Mouse α-Rat AF488 | Biolegend | RRID: AB_2910464<br>Cat#: 407513 |
| Mouse α-Rat AF594 | Biolegend | RRID: AB_2650845<br>Cat#: 407509 |
| Mouse α-Rabbit AF594 | Biolegend | RRID: AB_2832788<br>Cat#: 410407 |
| TruStain FcX™ Antibody | Biolegend | RRID: AB_1574973<br>Cat#: 101319 |
| α-CD45.1 BV785 | Biolegend | RRID: AB_2563379<br>Cat#: 110743 |
| α-CD45.1 PerCp-Cy5.5 | Biolegend | RRID: AB_893348<br>Cat#: 110727 |
| α-CD45.2 Pacific Blue | Biolegend | RRID: AB_492873<br>Cat#: 109819 |
| α-CD3 AF488 | Biolegend | RRID: AB_493530<br>Cat#: 100212 |
| α-CD3 PE | Biolegend | RRID: AB_312662<br>Cat#: 100205 |
| α-CD3 AF700 | Biolegend | RRID: AB_493696<br>Cat#: 100216 |
| α-CD4 BV421 | Biolegend | RRID: AB_11219790<br>Cat#: 100544 |
| α-CD4 PE | Biolegend | RRID: AB_313690<br>Cat#: 116005 |
| α-CD4 AF700 | Biolegend | RRID: AB_493698<br>Cat#: 100430 |
| α-CD8a BV510 | Biolegend | RRID: AB_2563057<br>Cat#: 100752 |
| α-CD11b PE | Biolegend | RRID: AB_312790<br>Cat#: 101207 |
| α-CD11b BV605 | Biolegend | RRID: AB_11126744<br>Cat#: 101237 |
| α-CD11c AF700 | Biolegend | RRID: AB_528735<br>Cat#: 117319 |
| α-CD19 PE-Cy7 | Biolegend | RRID: AB_313654<br>Cat#: 115519 |

| | | |
|---|---|---|
| α-CD25 PE-Cy5 | Biolegend | RRID: AB_312859<br>Cat#: 102010 |
| α-CD44 BV605 | Biolegend | RRID: AB_2562451<br>Cat#:103047 |
| α-CD49b APC | Biolegend | RRID: AB_313416<br>Cat#: 108909 |
| α-CD73 PE-Dazzle 594 | Biolegend | RRID: AB_2800628<br>Cat#: 127234 |
| α-CD172a APC | Biolegend | RRID: AB_2564060<br>Cat#: 144013 |
| α-CD274 PE-Dazzle594 | Biolegend | RRID: AB_2565638<br>Cat#: 124324 |
| α-CD278 BV650 | Biolegend | RRID: AB_2749928<br>Cat#: 313550 |
| α-CD279 BV750 | Biolegend | RRID: AB_2941421<br>Cat#: 135263 |
| α-CD304 PE | Biolegend | RRID: AB_2561927<br>Cat#: 145203 |
| α-CD317 PE | Biolegend | RRID: AB_1953284<br>Cat#: 127009 |
| α-B220 FITC | Biolegend | RRID: AB_312990<br>Cat#: 103206 |
| α-B220 PE | Biolegend | RRID: AB_312992<br>Cat#: 103207 |
| α-FR4 PE-Cy7 | Biolegend | RRID: AB_1134199<br>Cat#: 125012 |
| α-Gr-1 PE | Biolegend | RRID: AB_313372<br>Cat#: 108407 |
| α-Ter-119 PE | Biolegend | RRID: AB_313708<br>Cat#: 116207 |
| α-FOXP3 AF647 | Biolegend | RRID: AB_439749<br>Cat#: 320013 |
| α-Helios AF488 | Biolegend | RRID: AB_10645334<br>Cat#: 137213 |
| α-V$\beta$8.1 FITC | Biolegend | RRID: AB_1227787<br>Cat#: 118406 |
| α-V$\beta$11 PE | Biolegend | RRID: AB_10612759<br>Cat#: 139004 |
| α-CD117 APC | eBioscience | RRID: AB_469430<br>Cat#: 17-1171-82 |
| α-Sca-1 PE-Cy7 | eBioscience | RRID: AB_469669<br>Cat#: 25-5981-82 |
| α-CD117 | BioXCell | RRID: AB_2687818<br>Cat#: BE0293 |
| α-CD4 | BioXCell | RRID: AB_1107636<br>Cat#: BE0003-1 |
| α-CD8 | BioXCell | RRID: AB_10950145<br>Cat#: BE0117 |

| | | |
|---|---|---|
| **Chemicals, Peptides, and Recombinant Proteins** | | |
| LIVE/DEAD™ Fixable Near-IR Dead Cell Stain Kit | ThermoFisher | Cat#: L34975 |
| Bovine Serum Albumin | Fisher Scientific | Cat#: BP1600-100 |
| RBC Lysis Buffer (10X) | Biolegend | Cat#: 420301 |
| Cell Staining Buffer | Biolegend | Cat#: 420201 |
| True-Nuclear™ Transcription Factor Buffer Set | Biolegend | Cat#: 424401 |
| Propidium Iodide | MilliporeSigma | Cat#: P4170 |
| Liberase™ TL Research Grade | MilliporeSigma | Cat#: 05401020001 |
| Diphenhydramine HCl | Cayman Chemical Company | Cat#:11158 |
| Lineage Cell Depletion Kit, mouse | Miltenyi Biotec | Cat#:130-090-858 |
| HBS | Caisson Labs | Cat#: HBL06 |
| HEPES Solution | Caisson Labs | Cat#: HOL06 |
| RPMI 1640 | Corning | Cat#: 10-040-CV |
| LinBit | LinShin Canada, Inc | Cat#: Pr-1-B |
| LANTUS® | sanofi-aventis U.S. LLC | NDC 0088-2220-33 |
| UltraCruz® Hard-set Mounting Medium | Santa Cruz Biotechnology | Cat#: sc-359850 |
| Fetal Bovine Serum | Cytiva | Cat#: SH30070.03 |
| Penicillin-Streptomycin | Gibco | Cat#: 15140122 |
| **Experimental Models: Organisms/Strains** | | |
| B6 CD45.1 mice | The Jackson Laboratory | Stock #: 002014 |
| BALB/c mice | The Jackson Laboratory | Stock #: 000651 |
| FVB mice | The Jackson Laboratory | Stock #: 001800 |
| B6 *RIP-DTR* mice | Seung Kim Lab, Stanford University | N/A |
| **Software and Algorithms** | | |
| FlowJo 10.7 | FlowJo, LLC | N/A |
| GraphPad Prism 10 | GraphPad Software | N/A |
| Fiji | Ref 72. | N/A |
| **Other** | | |
| IC-250 X-Ray Biological Irradiator System | KIMTRON Inc | N/A |
| CM3050 S | Leica Biosystems | N/A |
| 5L Aurora System | Cytek | N/A |
| EVOS M5000 Cell Imaging System | ThermoScientific | N/A |
